# Hantavirus mutagenesis is impacted by mutagens and host antiviral proteins

**DOI:** 10.64898/2026.09.10.750593

**Authors:** Christopher Ruis, Julian Parkhill, R. Andres Floto

**Author notes:** Correspondence to: Christopher Ruis.

## Abstract

Hantaviruses are emerging zoonotic viruses that infect humans with high fatality rates. We currently have a poor understanding of the processes that contribute to hantavirus mutagenesis and therefore the potential for viral adaptation. To understand the drivers of mutations in hantaviruses, we calculated and compared mutational spectra across 14 divergent hantavirus species. We found that mutagens that are conserved across host species have a major impact on hantaviruses mutagenesis, contributing to at least six of the 12 possible mutation types. We found differences in mutational signatures across hantaviruses, some of which may be due to variation in host mutagen levels that are known to influence somatic mutations. However, we could find no evidence for differential mutational signatures between viruses associated with distinct diseases in humans and no evidence of tissue tropism differences between reservoir hosts. Using a novel method to quantify the impact of ZAP-mediated selection, we found that ZAP greatly influences mutational patterns in transition mutation types across hantaviruses. We also find evidence that APOBEC family proteins drive mutagenesis in all hantaviruses, with the likely exception of Nova virus. Our results therefore provide strong support that mutagens and host antiviral proteins are major contributors to hantavirus mutagenesis, thereby generating the genetic diversity necessary for viral adaptation.

## Introduction

Hantaviruses are emerging zoonotic viruses that naturally infect hosts within the orders Rodentia, Eulipotyphla and Chiroptera [1–3]. Each hantavirus typically exhibits a close association with one or a small number of host species within which it establishes predominantly asymptomatic infections [3]. For example, Hantaan virus lives within the striped field mouse (*Apodemus agrarius*), while Andes virus is carried by the long-tailed pygmy rice rat (*Oligozomys longicaudatus*) [2, 4]. This host association results in most hantaviruses exhibiting strong geographical restriction, determined by the range of their host [1, 3]. The exception is Seoul virus which infects black rats (*Rattus rattus)* and brown rats (*Rattus norvegicus*) and correspondingly exhibits a global distribution concomitant with that of its hosts [2].

Hantaviruses are capable of infecting multiple cell types, including endothelial cells, epithelial cells, macrophages, follicular dendritic cells and lymphocytes and can be detected in multiple tissues within their reservoir hosts including the lungs and kidneys [3, 5]. Within humans, hantaviruses can cause two distinct diseases, which are associated with different viruses and therefore geographical regions [1, 3, 6, 7]. Human hantavirus cases in Eurasia are associated with Hemorrhagic Fever with Renal Syndrome (HFRS), while those in the Americas are associated with Hantavirus Pulmonary Syndrome (HPS; also called Hantavirus Cardiopulmonary Syndrome or HCPS) [6, 7].

Human hantavirus infections predominantly occur through inhalation of secretions from the major host species, including faeces, saliva and urine [6, 7]. More rarely, infections can occur through bites [6]. Andes virus is currently the only hantavirus with documented human-to-human transmission; such events are associated with close contact and can lead to small clusters of cases [8, 9]. However, zoonotic transmission events are frequent enough to lead to an estimated 10,000 to 100,000 human hantavirus cases each year with an estimated fatality rate of <1% to 40% depending on the virus and outbreak [6].

Hantaviruses are within the genus *Orthohantavirus* and the Bunyaviridae family and as such have a negative sense single stranded RNA genome [10, 11]. The genome is organised into three segments, termed large (L), medium (M) and small (S), of which the M segment encodes the viral glycoproteins Gn and Gc [10]. The substitution rate of hantaviruses is estimated to be in the range of 10^-4^ to 10^-2^ substitutions/site/year, typical of RNA viruses [12]. This high substitution rate suggests that hantaviruses are capable of rapid adaptation.

Adaptation relies on the generation of diversity and therefore on mutagenesis. To understand microbial adaptation, we therefore need an understanding of the mutational processes that contribute to mutagenesis. However, we currently have a poor understanding of these factors within hantaviruses and therefore the processes that may assist or constrain hantavirus adaptation. We and others have recently demonstrated that mutational signatures (context-specific patterns of nucleotide substitution initially applied to detect the drivers of tumourigenesis [13–16]) can identify active mutational processes in microbes [17–26]. We have applied these methods to identify both intrinsic mutational processes encoded by the microbe and extrinsic lifestyle-associated mutagen exposures [18–20, 22]. In this study, we examine and compare mutational patterns across hantaviruses to identify mutational processes that may contribute to hantavirus adaptation.

## Results

### Mutagens impact hantavirus mutagenesis

We calculated single base substitution (SBS) mutational spectra for 14 hantavirus species that exhibit at least 600 sampled mutations [27] amongst M segment genetic sequences (***Figure 1, Table S1***, see ***Methods***). We additionally carried out “tree rescaling” of each spectrum by sequence composition across the respective phylogenetic tree [22]. The overall mutational spectrum is highly similar across viruses at the overall level (median cosine similarity 0.978; ***Figure S1***), supporting conservation of most mutational processes. We found that, as expected, each spectrum is dominated by transition mutations (median 76.7% unscaled, 78.1% tree rescaled; ***Figures 1, S2***). In contrast to SARS-CoV-2 [19, 26] and influenza A virus [22], T>A and A>T are typically the most common transversion mutations by raw count (***Figure S2***). However, following tree rescaling, G>T mutations occur at a similar rate to T>A and A>T mutations (***Figure S2***).

**Figure 1.**
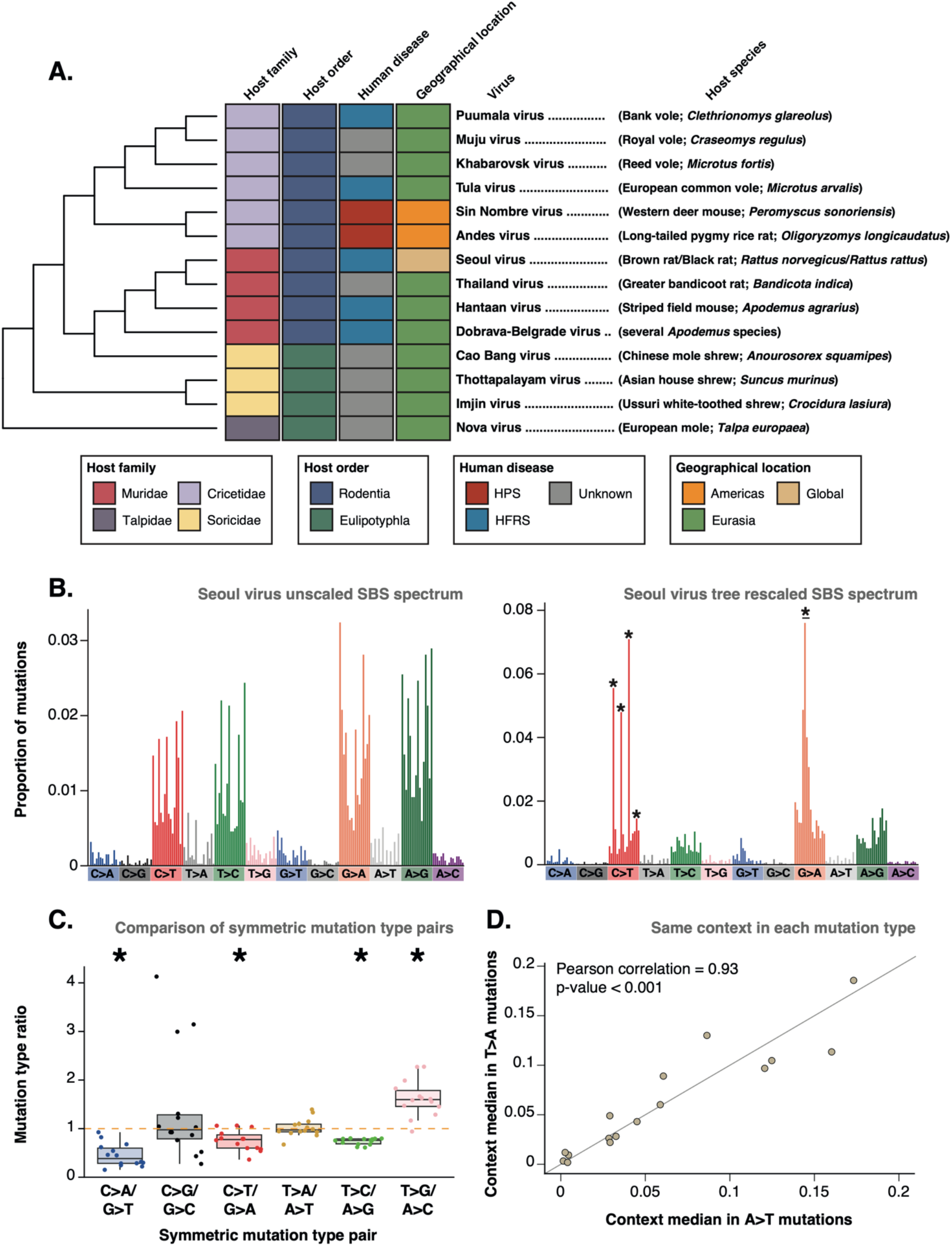
Mutagens influence hantavirus mutagenesis. (**A**) Cladogram of the evolutionary relationships between the 14 analysed hantaviruses. The heatmap shows the respective host family, host order, associated human disease (where known) and geographical distribution of each virus. Tip labels include the virus name and, in parentheses, the common English name and scientific name of the respective predominant host species. (**B**) The SBS mutational spectrum of Seoul virus is shown as proportions of raw mutation counts (left hand panel) and as the proportion of mutations within the tree rescaled spectrum (right hand panel). Asterisks in the right hand panel show C>T and G>A mutations that remove ZAP-associated CG contexts. (**C**) Comparison of symmetric mutation types in tree rescaled mutational spectra. * shows randomisation *P* < 0.05, supporting a significant difference from a ratio of one and therefore an elevation of one mutation type over the other. This supports a divergence from equal rates that would be expected under polymerase errors alone, and therefore supports the action of one or mutagens. (**D**) Comparison of contextual patterns between A>T and T>A mutations in tree rescaled mutational spectra. Each point shows the same mutational context in each mutation type, for example A[A>T]C and A[T>A]C. Points show median values across the eight hantaviruses with at least 160 A>T mutations and at least 160 T>A mutations; this filtering is applied to ensure there are sufficient mutations to compare contexts [22]. The significant positive correlation between non-symmetric contexts supports the action of one or more mutagens that causes A>T and T>A mutations with the same surrounding context preferences.

We next tested for the impact of mutagens by comparing the relative rates of symmetric mutation types [22] and found evidence of mutagens causing G>T (median 2.6-fold elevation), G>A (median 1.3-fold elevation), A>G (median 1.3-fold elevation) and T>G (median 1.6-fold elevation) mutations (permutation test *P* < 0.05, ***Figure 1***). We previously found evidence of mutagens causing G>T, A>G and T>G mutations in influenza A virus [22], which indicates either common mutagen exposures between the viruses or exposure to distinct sets of mutagens that induce similar mutational patterns. Conversely, we did not find evidence of a mutagen causing G>A mutations in influenza A virus [22], which indicates that hantaviruses are exposed to at least one mutagen that does not influence influenza A virus.

As T>A and A>T mutations occur at roughly equal rates after tree rescaling (***Figure 1, Figure S2***), their elevation relative to SARS-CoV-2 and influenza A virus may be due to either the hantavirus polymerase making more errors between A and T nucleotides, and/or elevated exposure to mutagens that cause A>T and T>A mutations. To distinguish between these possibilities, we examined the contextual preferences within these mutation types. If polymerase errors are responsible, A>T and T>A would exhibit symmetric contextual preferences [19, 22]. However, we observe only a weak correlation between symmetric contexts (***Figure S3***). Conversely, we observe a strong correlation between the same contexts in A>T and T>A mutations (for example, A[A>T]C and A[T>A]C; Pearson’s R = 0.93, randomisation *P* < 0.001; ***Figure 1***). Both mutations typically occur most frequently in CXG contexts (where X represents the mutated base) and occur very rarely in XT contexts (***Figure S4***). This strongly supports exposure to one or more mutagens that introduce A>T and T>A mutations at similar rates with the same contextual preferences. This further suggests that the elevation of A>T and T>A mutations in hantaviruses compared with influenza A virus and SARS-CoV-2 is due to mutagen exposure rather than polymerase errors.

### Drivers of differences in mutational patterns across hantaviruses

We next examined drivers of variation in mutational patterns between hantaviruses. We initially tested for an impact of viral evolutionary history and found no evidence that the proportion of any mutation type follows the hantavirus phylogenetic tree (continuous association index *P* > 0.05 in each case; ***Figure 2***). This indicates that viral genetic factors have limited direct impact on the mutational spectrum and therefore suggests that variation between viruses predominantly arises from extrinsic factors.

**Figure 2.**
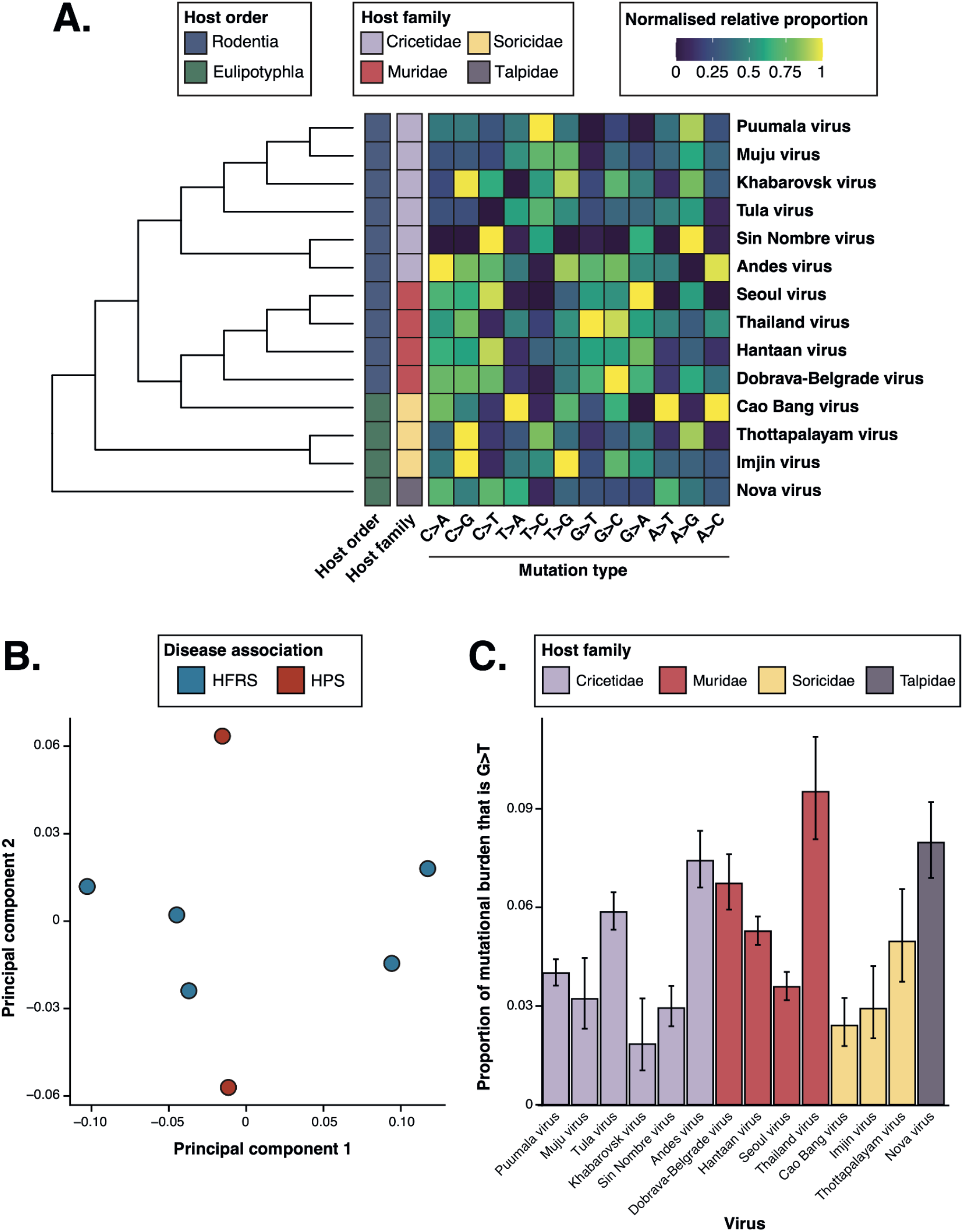
Variation in mutational spectrum across hantaviruses. (**A**) The cladogram shows the phylogenetic relationships between the 14 analysed hantaviruses [2]. Mutation types are shown as proportions within the respective spectrum and proportions are normalised so each column runs from zero to one. (**B**) Principal component analysis of seven hantaviruses with known human disease associations (***Table S1***) based on mutation type proportions. Viruses do not cluster by associated human disease. (**C**) The proportion of G>T mutations within the tree rescaled spectrum is shown for each hantavirus.

We further found no evidence that differences in mutational spectra are associated with host order or host family (ANOVA *P* > 0.05, ***Figure S5***), or that any mutation type follows the host phylogeny (continuous association index *P* > 0.05, ***Table S2***). This suggests that the hantavirus mutational spectrum is not greatly influenced by host factors that are conserved at the level of host family or host order.

We next tested whether there is evidence of differential mutational spectra between hantaviruses associated with distinct disease in humans. We found no separation of viruses associated with HFRS from those associated with HPS when clustering based on mutational patterns and observed no mutation types that differ by associated disease (ANOVA *P* > 0.05; ***Figures 2, S6***). This suggests that differences in human disease between viruses are not driven by alternative tissue tropism within the reservoir species.

We next examined variation in the level of G>T mutations across hantaviruses, which we have previously found can be associated with replication sites in RNA viruses and bacteria [18, 19]. We found that the level of G>T mutations varies markedly between hantaviruses, from 1.8% of mutational burden (accounting for genomic composition) in Khabarovsk virus to 9.5% in Thailand virus (***Figure 2***). This variation may be driven by differential replication site preferences in different viruses and/or by distinct mutagen levels across host species. In potential support of the latter scenario, Seoul virus exhibits lower G>T mutations than the other murinae-associated hantaviruses (***Figure 2***). Previous work has found a lower rate of somatic G>T mutations in gastrointestinal tissue of *Rattus norvegicus* (a major natural host of Seoul virus [2]) compared to *Mus musculus* [28]. This may suggest that mutagens produced by the host can impact the genetic material of both host and virus and further indicate that variation in G>T level across hantaviruses is influenced by the level of one or more mutagens produced by the host.

### Impact of host antiviral proteins on hantavirus mutagenesis

We next examined whether host antiviral proteins may impact hantavirus mutational patterns. We initially developed a “ZAP enrichment score” that calculates the relative ZAP activity by comparing contexts impacted by ZAP with those not impacted by ZAP (see ***Methods***). This score is applied to each mutation type within the tree rescaled mutational spectrum of each virus and examines evidence for ZAP activity through the magnitude of the enrichment (for mutation types that remove ZAP contexts) or suppression (for mutation types that introduce ZAP contexts) of ZAP-associated contexts.

We observe consistent evidence of ZAP-mediated selection in C>T, T>C and A>G mutations (permutation test *P* < 0.05; ***Figures 1, 3***). However, we do not observe evidence of reduction of ZAP-associated mutations in T>G mutations, similar to influenza A virus [22]. This indicates that ZAP-mediated selection predominantly acts upon high frequency transition mutations. Alternatively, as T>G is caused by at least one mutagen (***Figure 1***), ZAP-mediated selection may be outweighed by this mutagen. While several hantaviruses exhibit an elevation of G>A mutations in ZAP-associated contexts, we do not observe a consistent elevation (***Figure 3***). In particular, Cao Bang virus is an outlier within G>A mutations and does not contain any sampled G>A mutations in ZAP-associated contexts (***Figure S7***), despite 190 sampled G>A mutations. Cao Bang virus is not an outlier for other mutation types, suggesting that this is not due to reduced ZAP-mediated selection. We find a significant correlation between the magnitude of ZAP enrichment in G>A mutations and the C>T/G>A symmetric mutation type ratio (permutation test *P* < 0.05 including and excluding Cao Bang virus; ***Figure S8***). This may indicate that hantaviruses with reduced ZAP enrichment in G>A mutations are more exposed to an additional mutagen driving G>A mutations under different contextual preferences, which reduces the overall enrichment of ZAP-associated contexts.

**Figure 3.**
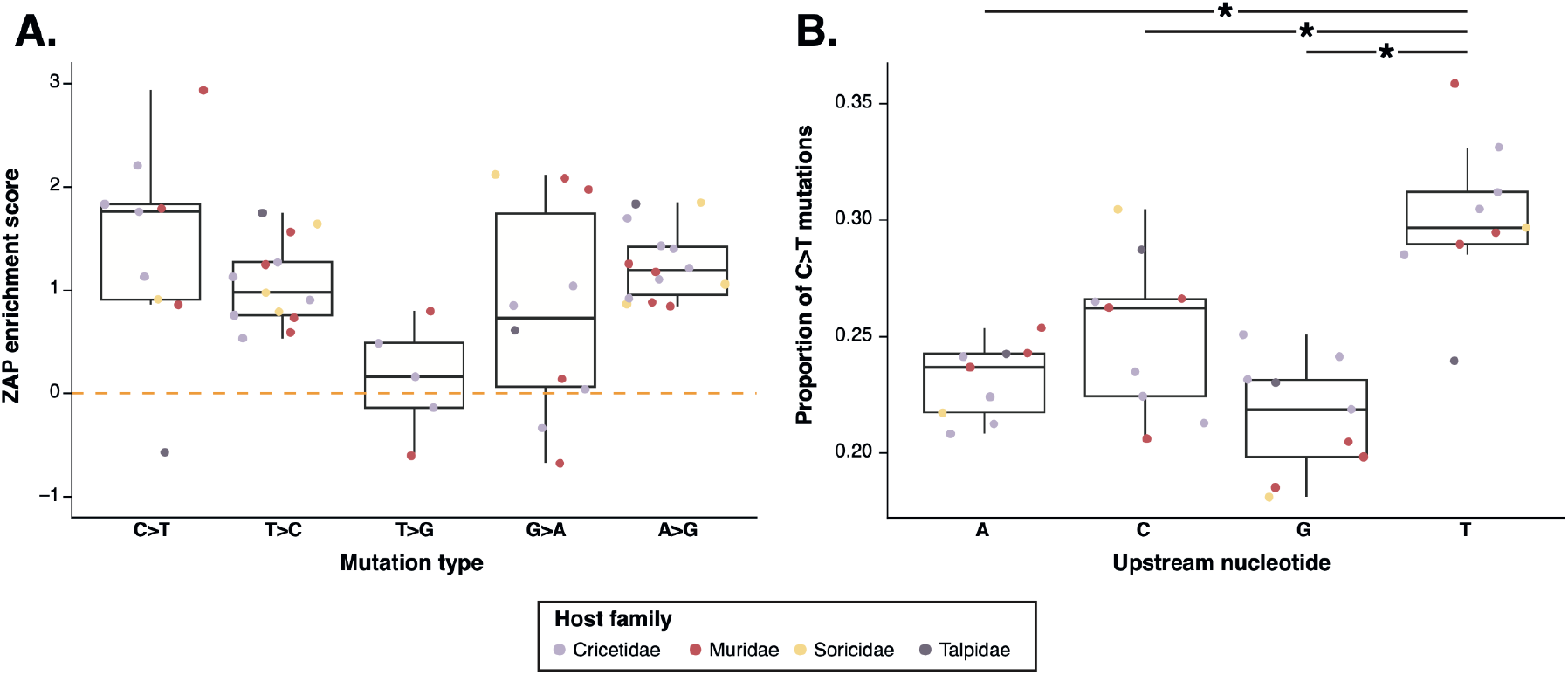
Influence of host ZAP and APOBEC family proteins on hantavirus mutagenesis. (**A**) We developed a ZAP enrichment score that calculates relative ZAP activity through the enrichment (for mutation types that remove ZAP contexts) or suppression (for mutation types that introduce ZAP contexts) of ZAP-associated contexts. This score was calculated for each mutation type in each virus that contains at least 160 mutations within the respective mutation type (see ***Methods***). The distributions of these scores are shown for each mutation type. T>A and A>T mutations are not included as they do not introduce or remove ZAP contexts. C>A, C>G, G>C, G>T and A>C are not shown as they contain a maximum of two viruses with at least 160 mutations. (**B**) The proportion of C>T mutations with each upstream nucleotide in tree rescaled spectra. * shows permutation test *P* < 0.05 comparing the distributions of T[C>T] mutations with those of the other upstream nucleotides.

We do not find evidence that any hantavirus exhibits consistently higher ZAP activity across mutation types, or of consistent elevation of ZAP activity within a host family (***Figure 2***). This suggests that ZAP activation and activity is similar across hosts of the examined hantaviruses.

While ZAP-mediated selection has a major influence of contextual preferences, we find evidence of additional contextual preferences across mutation types that are not attributable to ZAP (***Figure S4***). This supports the action of additional context-specific mutational processes.

Comparison of contextual patterns within C>T mutations showed elevation of mutations in T[C>T] contexts over each other nucleotide context (permutation test *P* < 0.05, ***Figure 2***), consistent with the action of APOBEC-induced mutagenesis [21, 23]. However, we found lower T[C>T] mutations within Nova virus compared to other hantaviruses (more than 2.5 median absolute deviations away from the median; ***Figure 2***). While the genome of the European mole (*Talpa europaea*; the host of Nova virus) contains four annotated APOBEC genes (***Table S3***), the same number as other hantavirus hosts with available genomic data, this may be indicative of reduced activity of one or more APOBEC proteins in this host, or elevated evasion of one or more APOBEC proteins by Nova virus.

Our results support the host antiviral proteins ZAP and APOBEC having a major influence on hantavirus mutational patterns.

## Discussion

Mutations enable viral adaptation and may be introduced by polymerase errors during replication or by exposure to external mutagenic processes. It is possible to distinguish these processes as polymerase errors introduce symmetric mutational patterns while external mutagenic processes typically cause asymmetric mutations. We found evidence of deviation from symmetric mutation type ratios and contexts in at least six of the 12 mutation types (***Figure 1***), supporting mutagen exposure, and enrichment of mutations in APOBEC-associated contexts and contextual preferences associated with ZAP-mediated selection (***Figure 3***), supporting the action of host antiviral proteins on viral evolution. Our results therefore provide strong support that hantavirus mutations are not introduced solely by the error prone viral polymerase.

Mutagen-induced patterns are present across analysed hantaviruses (***Figure 1***), which suggests that the responsible mutagens are present across their host species. We identified similarities between mutagen exposures in hantaviruses and influenza A virus [22], which may indicate general exposure of RNA viruses to specific core mutagens. While influenza A virus genome replication occurs within the cell nucleus [29], hantaviruses do not enter the nucleus during their replication cycle [10]. This suggests that these putative mutagens are likely to be present within the host cell cytoplasm rather than the nucleus. Future work will be needed to examine whether these viruses are exposed to the same set of mutagens or to distinct mutagens with equivalent impacts on mutation types.

Notably, hantaviruses are exposed to at least one mutagen causing G>A mutations (***Figure 1***). This mutagen either does not influence influenza A virus mutagenesis or influenza A virus is exposed to an additional mutagen that causes C>T mutations at a similar level [22]. In either scenario, this provides strong support for differential mutagen exposure between hantaviruses and influenza A virus. This could be due to differences in the viral replication cycle, differential activation of host pathways, and/or distinct mutagen production by different hosts.

We found no evidence for distinct mutational patterns in hantaviruses associated with different disease syndromes in humans (***Figures 2, S6***). As we have previously found strong evidence of differential mutational patterns between host body sites [18, 19, 22], this suggests that the type of human disease a hantavirus causes is not driven by differences in tissue tropism in the reservoir hosts. Rather, this is likely to be due to viral mutations maintained due to other processes that result in distinct interactions with human tissues.

Our results provide strong support that the host antiviral proteins ZAP and members of the APOBEC family impact hantavirus mutational patterns (***Figure 3***). We found consistent ZAP activity across viruses, indicating that the ZAP proteins of all hantavirus hosts are similarly active. While CpG suppression, indicating historical ZAP exposure, has previously been noted in hantaviruses [30], our results provide evidence of ZAP activity during recent hantavirus circulation. Together, our results provide strong support that mutagens and host antiviral proteins are major factors influencing hantavirus mutagenesis, and consequently ongoing evolution.

## Methods

### Calculation of hantavirus mutational spectra

We aimed to calculate mutational spectra for all hantaviruses with genomic data. We focussed our analyses on the M segment as this typically has the largest number of available genetic sequences. To obtain sequence datasets, we searched NCBI Genome with the name of the virus and downloaded all available M segment sequences. Sequences were filtered to retain only those that contain at least 80% of the length of the M segment within the respective virus reference sequence in NCBI genome. We excluded sequences associated with patents or methods development. The majority of known hantaviruses have only a small number of M segment sequences on NCBI GenBank; we retained 17 hantaviruses that have at least five M segment sequences for further analysis.

Sequences were aligned at the amino acid level using MUSCLE [31] in SEAVIEW v5.0.4 [32]. Alignments were manually inspected and adjusted as necessary within SEAVIEW [32]. We reconstructed a maximum likelihood phylogenetic tree for sequences within each virus using IQ-TREE v2.1.3 [33] employing the HKY model of nucleotide substitution with gamma rate heterogeneity and four gamma classes. Within each dataset, we included an outgroup sequence from a closely related hantavirus (identified based on phylogenetic trees in previous publications [2]) which was used to root the phylogenetic tree.

We calculated mutational spectra using MutTui v2.0.2. The mutational spectra of Puumala virus and Muju virus and for Cao Bang virus and Lianghe virus were calculated simultaneously using a single phylogenetic tree containing sequences from both viruses; phylogenetic branches were labelled by virus so mutational spectra were calculated separately.

Asama virus, Lianghe virus and Dabieshan virus contained fewer than 600 sampled mutations within their mutational spectrum (***Table S1***). These viruses were therefore excluded from further analyses [27]. We therefore obtained a final mutational spectrum dataset of 14 hantaviruses that contain at least 600 sampled mutations (***Table S1***).

All mutational spectra were additionally “tree rescaled” [22]. This rescales the SBS mutation counts by the sequence composition across the respective phylogenetic tree, thereby accounting for the opportunity for each SBS mutation to occur. We employed tree rescaled mutational spectra for analyses that could be impacted by sequence composition.

When examining SBS contextual patterns within a mutation type, we retained only viruses that have at least 160 sampled mutations within the mutation type, as previously [22].

### Assessment of mutagen activity

We examined mutagen activity by comparing the ratio of symmetric mutation type pairs in tree rescaled mutational spectra [22]. To identify symmetric mutation type pairs where one mutation type is significantly elevated over the other, we compared the median mutation ratio in real data with the distribution of median mutation ratios across 1000 randomisations of mutation type proportions across the mutation types in the symmetric mutation type pair. The p-value was calculated as the proportion of randomisations with a mutation type ratio at least as large as with the real data. Benjamini-Hochberg correction was applied to account for multiple testing.

### Comparison of contextual patterns in A>T and T>A mutations

To examine potential mechanisms for the elevation of A>T and T>A mutations, we compared symmetric mutational contexts and the same mutational contexts. We calculated the Pearson’s correlation coefficient between each set of contexts in A>T and T>A mutations. To assess statistical significance, we compared the correlation in the real data with the distribution of correlations in 1000 randomisations of T>A proportions across contexts. The p-value was calculated as the proportion of randomisations with a correlation at least as large as with the real data.

### Examination of the drivers of variation between hantavirus mutational spectra

To examine the impact of viral evolutionary history on mutational spectrum, we reconstructed a cladogram of the evolutionary relationships between the 14 analysed hantaviruses based on previous studies [2]. We then examined each mutation type independently. For each mutation type, each phylogenetic tip is labelled with the respective proportion of that mutation type and the continuous association index calculated [34, 35]. The significance of the association index is assessed by comparing the value in real data with the distribution of association indices across 1000 randomisations of proportions across tips. The p-value is the proportion of randomisations with a value at least as large as with the real data. Benjamini-Hochberg correction was applied to account for multiple testing. No mutation types exhibited statistical significance (*P* > 0.05).

We additionally tested for an impact of host phylogeny as above but applying the phylogenetic tree of host relationships rather than virus relationships. The phylogenetic tree of hosts was reconstructed based on previous genetic analyses of Rodentia and Eulipotyphla evolution [36, 37].

To test for an impact of host order or host family on mutational patterns, we carried out analysis of variance (ANOVA) comparing the respective host groups for each mutation type with Benjamini-Hochberg correction applied. We applied the same methodology to test for an impact of known human-associated disease, but here filtered the dataset to retain only the seven hantaviruses that have a known disease association (HFRS or HPS).

### Calculation of ZAP context enrichment

We developed a score to assess the degree of ZAP-associated context enrichment within each mutation type. This score is calculated within tree rescaled mutational spectra. We calculate the ZAP enrichment score within a mutation type using:

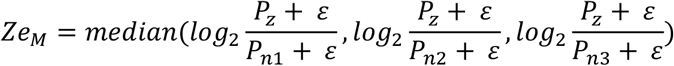

Where: *Ze* is the ZAP enrichment score; *M* is the mutation type being considered; *Pz* is the proportion of the mutational burden within mutation type *M* that is within ZAP-associated contexts; *Pn1, Pn2* and *Pn3* are the proportions of the mutational burden within mutation type *M* that is within each of the other three non-ZAP associated contexts; and ε is a small constant (0.000001) that is applied to account for the potential for zero mutations within a given context.

The contexts within *Pz, Pn1, Pn2* and *Pn3* depends on the mutation type. Where the mutation occurs at the first position of the CG dinucleotide, the context will be the downstream nucleotide. Where the mutation occurs at the second position of the CG dinucleotide, the context will be the upstream nucleotide.

ZAP-mediated selection is expected to increase mutations in mutation types that remove CG contexts; in these cases, ZAP activity will result in positive *Ze* scores. However, ZAP-mediated selection is expected to decrease mutations in mutation types that introduce CG contexts. To enable comparison across mutation types, we therefore take the absolute value of *Ze* for all mutation types. The score therefore takes into account the direction of the expected impact so a larger score in each mutation type is associated with elevated ZAP activity.

### Comparison of the magnitude of ZAP enrichment in G>A mutations and the C>T/G>A symmetric mutation type ratio

To examine the correlation between ZAP enrichment in G>A mutations and the C>T/G>A symmetric mutation type ratio, we calculated the Pearson’s correlation with the real data and compared this with the distribution of correlations in 1000 randomisations of the C>T/G>A ratio across datasets. The p-value was calculated as the proportion of randomisations with a correlation at least as large as with the real data. To prevent interpretation being influenced by the outlier Cao Bang virus, we carried out this analysis with and without Cao Bang virus included.

### Identification of over-represented and under-represented SBS contexts

We identified SBS mutation contexts that are over-represented or under-represented across tree rescaled mutational spectra as those whose median proportion across all spectra is more than 2.5 times the median absolute deviation away from the median across all contexts. This analysis was carried out on each mutation type independently, filtering to retain only those viruses with at least 160 mutations within the respective mutation type.

## Data summary

All sequence alignments, phylogenetic trees and mutational spectra are available at https://github.com/chrisruis/hantavirus_mutational_spectra.

## Author contributions

C.R. conceived the project. C.R. and R.A.F. designed the experiments. C.R., J.P. and R.A.F. wrote the manuscript. C.R. performed phylogenetic and mutational spectrum analyses. J.P. and R.A.F. provided supervisory support.

## References

1. Holmes EC, Zhang Y-Z. The evolution and emergence of hantaviruses. Current Opinion in Virology 2015;10:27–33.

2. Guo W-P, Lin X-D, Wang W, Tian J-H, Cong M-L, et al. Phylogeny and Origins of Hantaviruses Harbored by Bats, Insectivores, and Rodents. PLOS Pathogens 2013;9:e1003159.

3. Jonsson CB, Figueiredo LTM, Vapalahti O. A Global Perspective on Hantavirus Ecology, Epidemiology, and Disease. Clinical Microbiology Reviews 2010;23:412–441.

4. Jeyachandran AV, Irudayam JI, Dubey S, Chakravarty N, Daskou M, et al. Differential tropisms of old and new world hantaviruses influence virulence and developing host-directed antiviral candidates. PLoS Pathog 2025;21:e1013401.

5. Noack D, Goeijenbier M, Reusken CBEM, Koopmans MPG, Rockx BHG. Orthohantavirus Pathogenesis and Cell Tropism. Frontiers in Cellular and Infection Microbiology;10. https://www.frontiersin.org/articles/10.3389/fcimb.2020.00399 (2020, accessed 1 November 2022).

6. World Health Organization. Hantavirus. https://www.who.int/news-room/fact-sheets/detail/hantavirus.

7. Vial PA, Ferrés M, Vial C, Klingström J, Ahlm C, et al. Hantavirus in humans: a review of clinical aspects and management. The Lancet Infectious Diseases 2023;23:e371–e382.

8. Padula PJ, Edelstein A, Miguel SD, López NM, Rossi CM, et al. Hantavirus pulmonary syndrome outbreak in Argentina: molecular evidence for person-to-person transmission of Andes virus. Virology 1998;241:323–330.

9. Martinez-Valdebenito C, Calvo M, Vial C, Mansilla R, Marco C, et al. Person-to-Person Household and Nosocomial Transmission of Andes Hantavirus, Southern Chile, 2011. Emerg Infect Dis 2014;20:1629–1636.

10. Meier K, Thorkelsson SR, Quemin ERJ, Rosenthal M. Hantavirus Replication Cycle—An Updated Structural Virology Perspective. Viruses 2021;13:1561.

11. Muyangwa M, Martynova EV, Khaiboullina SF, Morzunov SP, Rizvanov AA. Hantaviral Proteins: Structure, Functions, and Role in Hantavirus Infection. Front Microbiol 2015;6:1326.

12. Ramsden C, Melo FL, Figueiredo LuizM, Holmes EC, Zanotto PMA, et al. High Rates of Molecular Evolution in Hantaviruses. Mol Biol Evol 2008;25:1488–1492.

13. Nik-Zainal S, Kucab JE, Morganella S, Glodzik D, Alexandrov LB, et al. The genome as a record of environmental exposure. Mutagenesis 2015;30:763–770.

14. Nik-Zainal S, Alexandrov LB, Wedge DC, Van Loo P, Greenman CD, et al. Mutational Processes Molding the Genomes of 21 Breast Cancers. Cell 2012;149:979–993.

15. Alexandrov LB, Nik-Zainal S, Wedge DC, Aparicio SAJR, Behjati S, et al. Signatures of mutational processes in human cancer. Nature 2013;500:415–421.

16. Alexandrov LB, Kim J, Haradhvala NJ, Huang MN, Tian Ng AW, et al. The repertoire of mutational signatures in human cancer. Nature 2020;578:94–101.

17. Ruis C, Bryant JM, Bell SC, Thomson R, Davidson RM, et al. Dissemination of Mycobacterium abscessus via global transmission networks. Nat Microbiol 2021;1–10.

18. Ruis C, Weimann A, Tonkin-Hill G, Pandurangan AP, Matuszewska M, et al. Mutational spectra are associated with bacterial niche. Nat Commun 2023;14:7091.

19. Ruis C, Peacock TP, Polo LM, Masone D, Alvarez MS, et al. A lung-specific mutational signature enables inference of viral and bacterial respiratory niche. Microbial Genomics 2023;9:001018.

20. Sanderson T, Hisner R, Donovan-Banfield I, Hartman H, Løchen A, et al. A molnupiravir-associated mutational signature in global SARS-CoV-2 genomes. Nature 2023;1–3.

21. Otieno JR, Ruis C, Onoja AB, Kuppalli K, Hoxha A, et al. Global genomic surveillance of monkeypox virus. Nat Med 2024;1–1.

22. Ruis C, Rossi AD, Sanderson T, Matuszewska M, Ntemourtsidou M, et al. Mutational spectra reveal influenza virus transmission routes and adaptation. 2025;2025.11.26.690773.

23. O’Toole Á, Neher RA, Ndodo N, Borges V, Gannon B, et al. APOBEC3 deaminase editing in mpox virus as evidence for sustained human transmission since at least 2016. Science 2023;382:595–600.

24. McBride DS, Garushyants SK, Franks J, Magee AF, Overend SH, et al. Accelerated evolution of SARS-CoV-2 in free-ranging white-tailed deer. Nat Commun 2023;14:5105.

25. Corbett-Detig R. A phylogenetic method identifies candidate drivers of the evolution of the SARS-CoV-2 mutation spectrum. Molecular Biology and Evolution 2025;msaf059.

26. Bloom JD, Beichman AC, Neher RA, Harris K. Evolution of the SARS-CoV-2 Mutational Spectrum. Mol Biol Evol 2023;40:msad085.

27. Ruis C, Tonkin-Hill G, Floto RA, Parkhill J. Calculating and applying pathogen mutational spectra using MutTui. 2023;2023.06.15.545111.

28. Cagan A, Baez-Ortega A, Brzozowska N, Abascal F, Coorens THH, et al. Somatic mutation rates scale with lifespan across mammals. Nature 2022;604:517–524.

29. Carter T, Iqbal M. The Influenza A Virus Replication Cycle: A Comprehensive Review. Viruses 2024;16:316.

30. Forni D, Pozzoli U, Cagliani R, Clerici M, Sironi M. Dinucleotide biases in RNA viruses that infect vertebrates or invertebrates. Microbiology Spectrum 2023;11:e02529–23.

31. Edgar RC. MUSCLE: a multiple sequence alignment method with reduced time and space complexity. BMC Bioinformatics 2004;5:113.

32. Gouy M, Guindon S, Gascuel O. SeaView Version 4: A Multiplatform Graphical User Interface for Sequence Alignment and Phylogenetic Tree Building. Molecular Biology and Evolution 2010;27:221–224.

33. Minh BQ, Schmidt HA, Chernomor O, Schrempf D, Woodhams MD, et al. IQ-TREE 2: New Models and Efficient Methods for Phylogenetic Inference in the Genomic Era. Molecular Biology and Evolution 2020;37:1530–1534.

34. Wang TH, Donaldson YK, Brettle RP, Bell JE, Simmonds P. Identification of Shared Populations of Human Immunodeficiency Virus Type 1 Infecting Microglia and Tissue Macrophages outside the Central Nervous System. J Virol 2001;75:11686–11699.

35. Parker J, Rambaut A, Pybus OG. Correlating viral phenotypes with phylogeny: accounting for phylogenetic uncertainty. Infect Genet Evol 2007;8:239–246.

36. Steppan SJ, Schenk JJ. Muroid rodent phylogenetics: 900-species tree reveals increasing diversification rates. PLoS One 2017;12:e0183070.

37. Yuan H, Dickson ED III, Martinez Q, Arnold P, Asher RJ. The origin and evolution of shrews (Soricidae, Mammalia). Proc R Soc B 2024;291:20241856.

