## Supplementary figures for "Hantavirus mutagenesis is impacted by mutagens and host antiviral proteins"

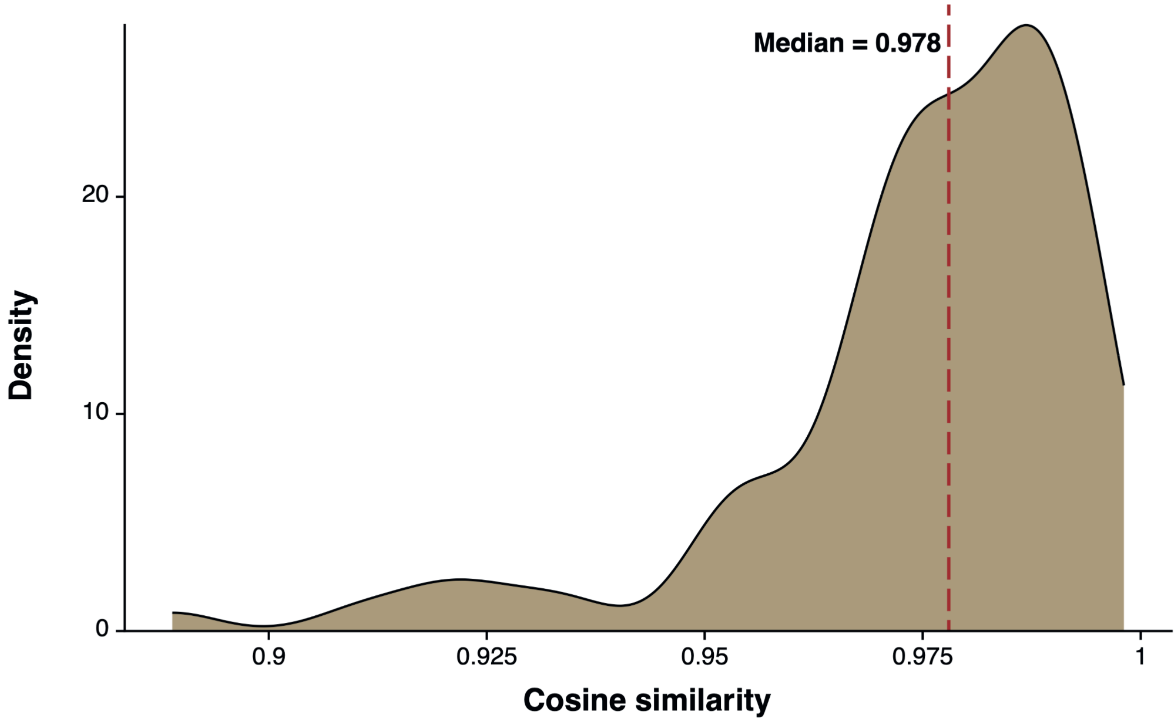


**Figure S1. Hantavirus mutational spectra are similar at the overall level.** The distribution of cosine similarities between all pairs of hantavirus SBS mutational spectra is shown.

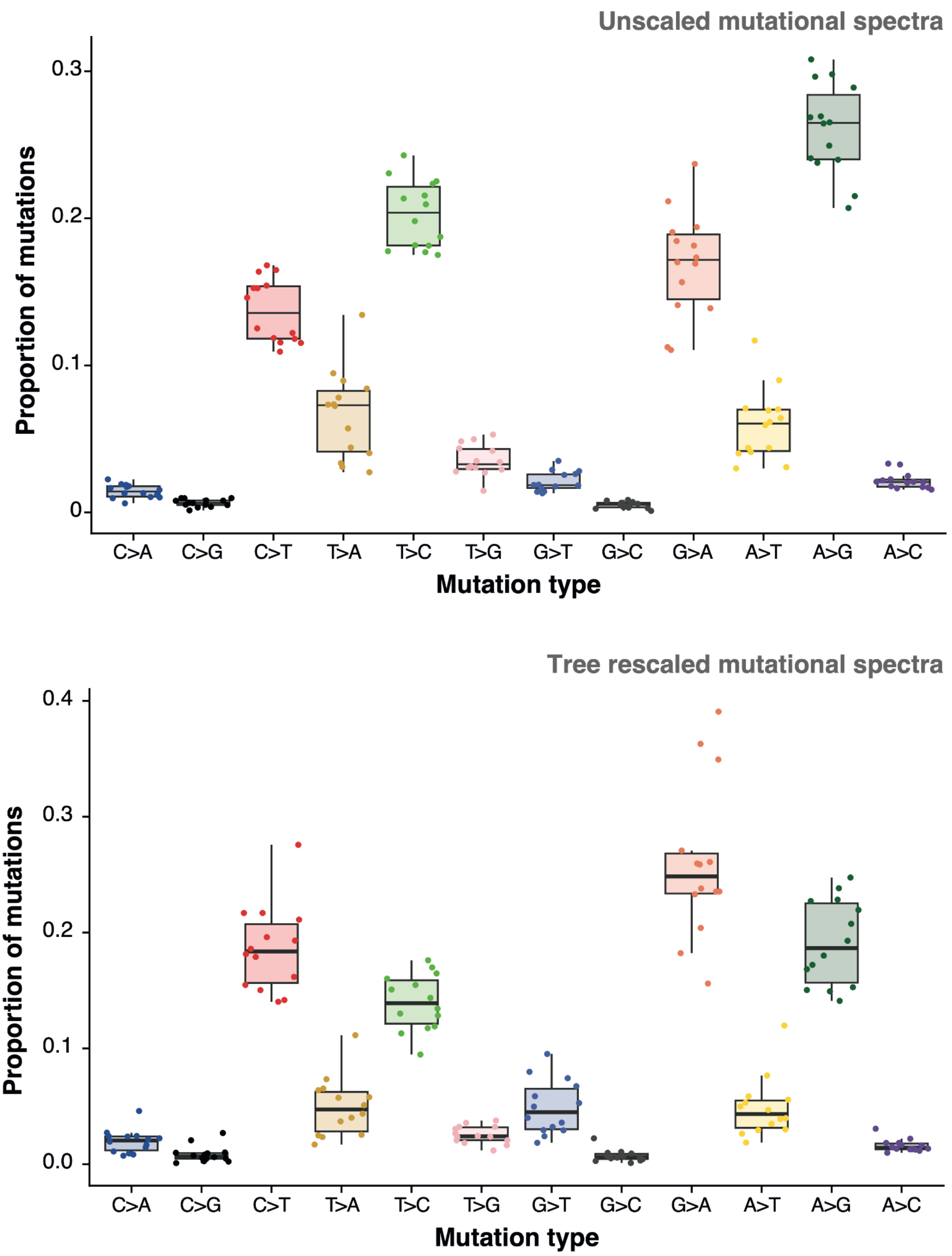


**Figure S2. Mutation type composition of hantavirus mutational spectra.** The proportion of each mutation type is shown for each hantavirus mutational spectrum in unscaled mutational spectra (**top panel**) and tree rescaled mutational spectra (**bottom panel**).

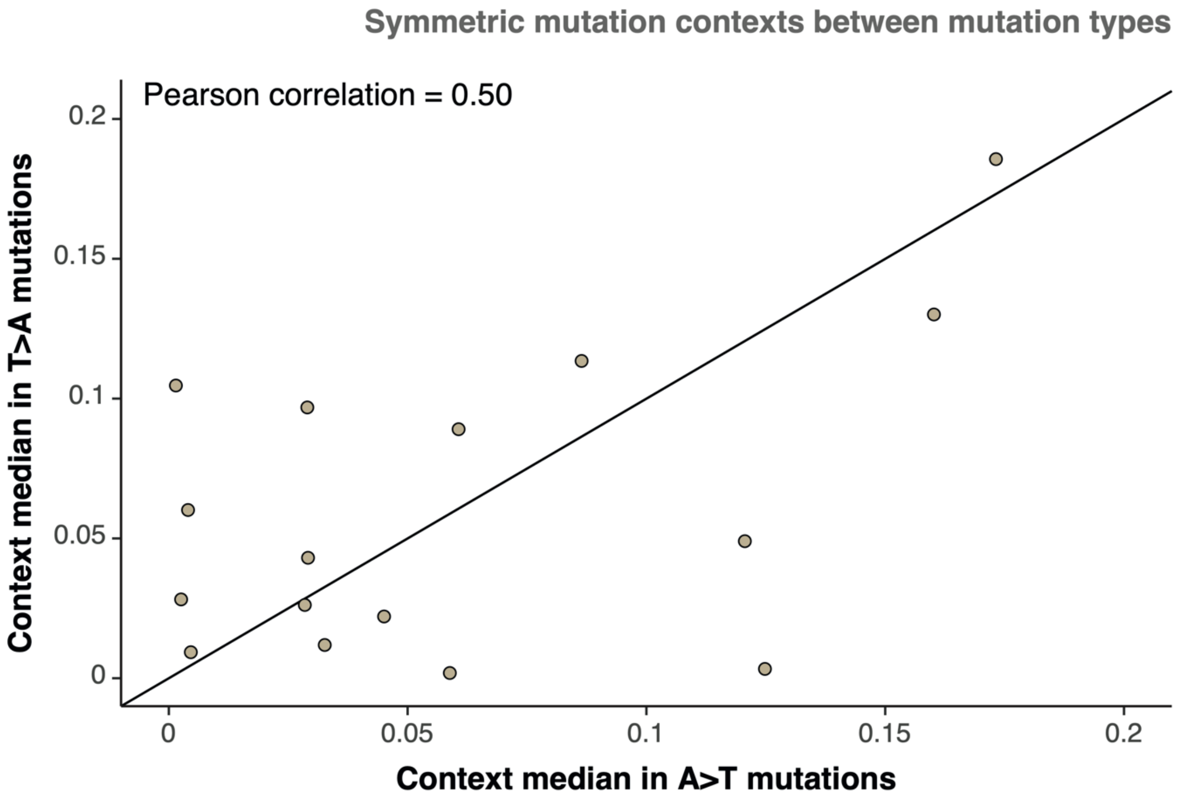


**Figure S3. Comparison of symmetric contexts in A>T and T>A mutations across hantaviruses.** Each point represents the comparison between the symmetric SBS contexts in A>T and T>A mutations, for example A[A>T]C and G[T>A]T. Median values across the eight hantaviruses with at least 160 A>T mutations and at least 160 T>A mutations are shown. This filtering is applied to ensure there are sufficient mutations within each mutation type to compare. Comparisons are carried out using tree rescaled mutational spectra.

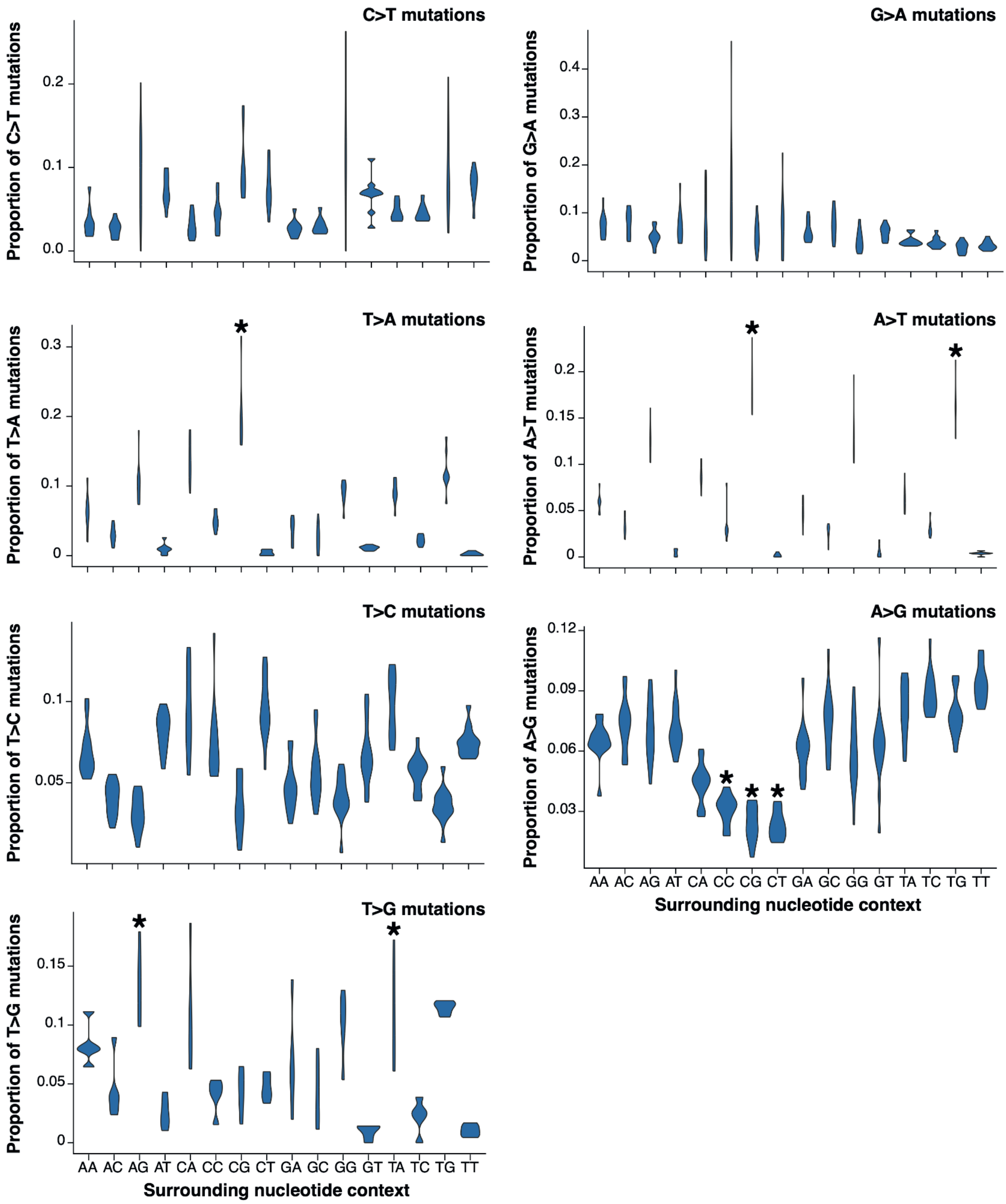


**FIgure S4. Contextual preferences in hantavirus mutational patterns.** For each mutation type, we filtered to include hantaviruses where the respective mutation type has at least 160 mutations. The proportion of mutations within the respective mutation type that occur within each context is shown. * shows mutation contexts identified as significantly elevated or reduced, identified as those with median proportion more than 2.5 times the median absolute deviation away from the median context proportion for the mutation type.

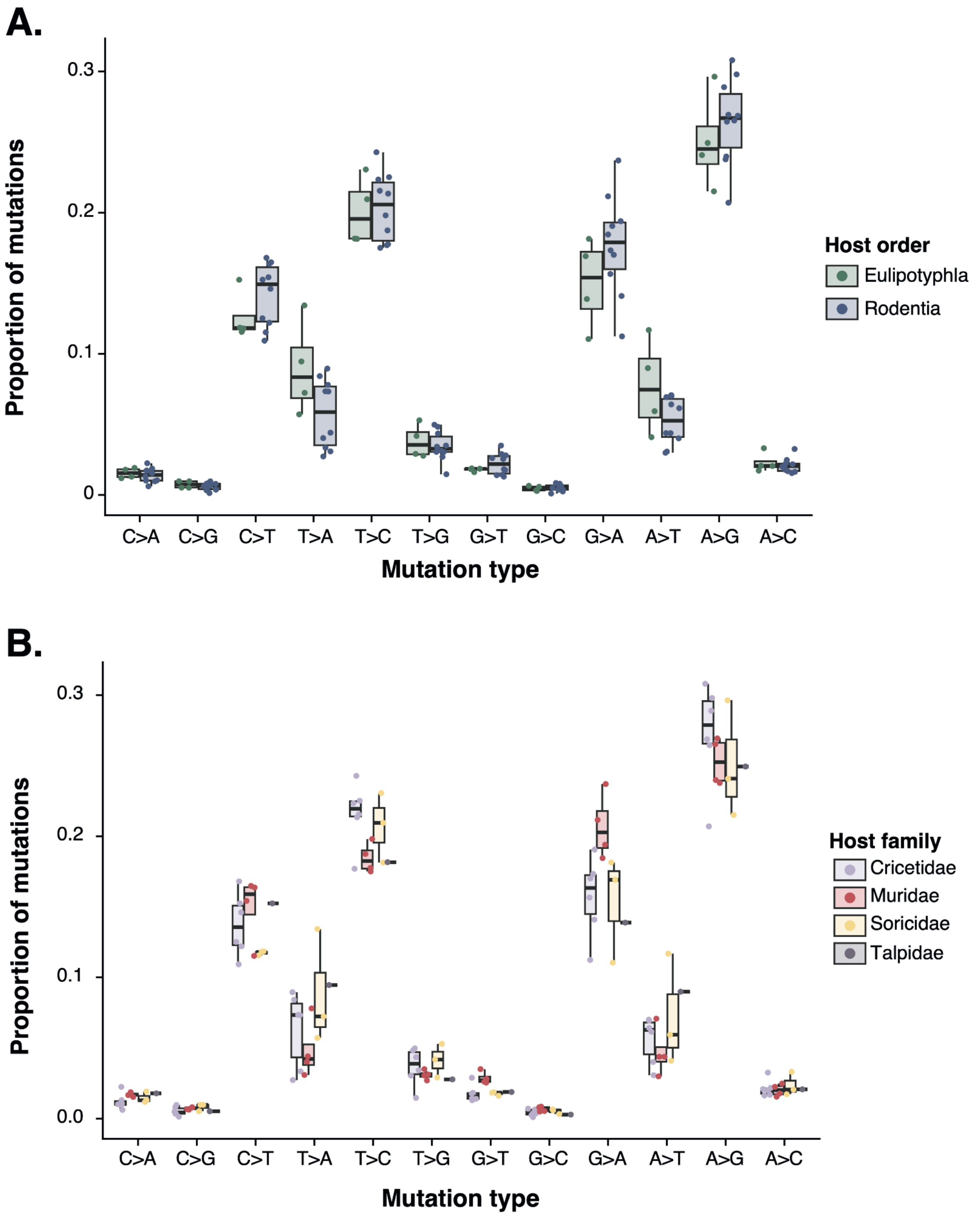


**Figure S5. No mutation types differ significantly between host orders or host families.** The proportion of each mutation type in each hantavirus spectrum is shown separated by (**A**) the taxonomic order of the host species and (**B**) the host family. No mutation type differs significantly by host order or host family (Benjamini Hochberg-adjusted ANOVA p-value > 0.05).

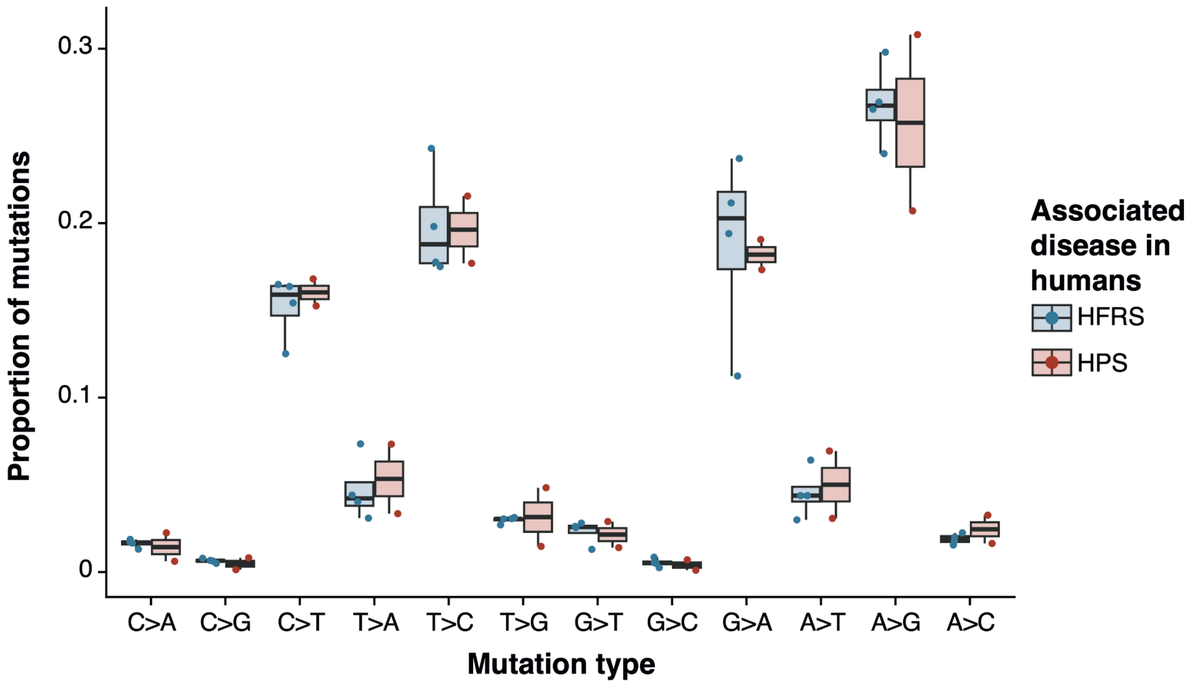


**Figure S6. No mutation types differ by human-associated disease.** We filtered to retain the seven hantaviruses that have a known disease association in humans. The proportion of each mutation type is shown for each hantavirus. No mutation type differs significantly by host order or host family (Benjamini Hochberg-adjusted ANOVA p-value > 0.05).

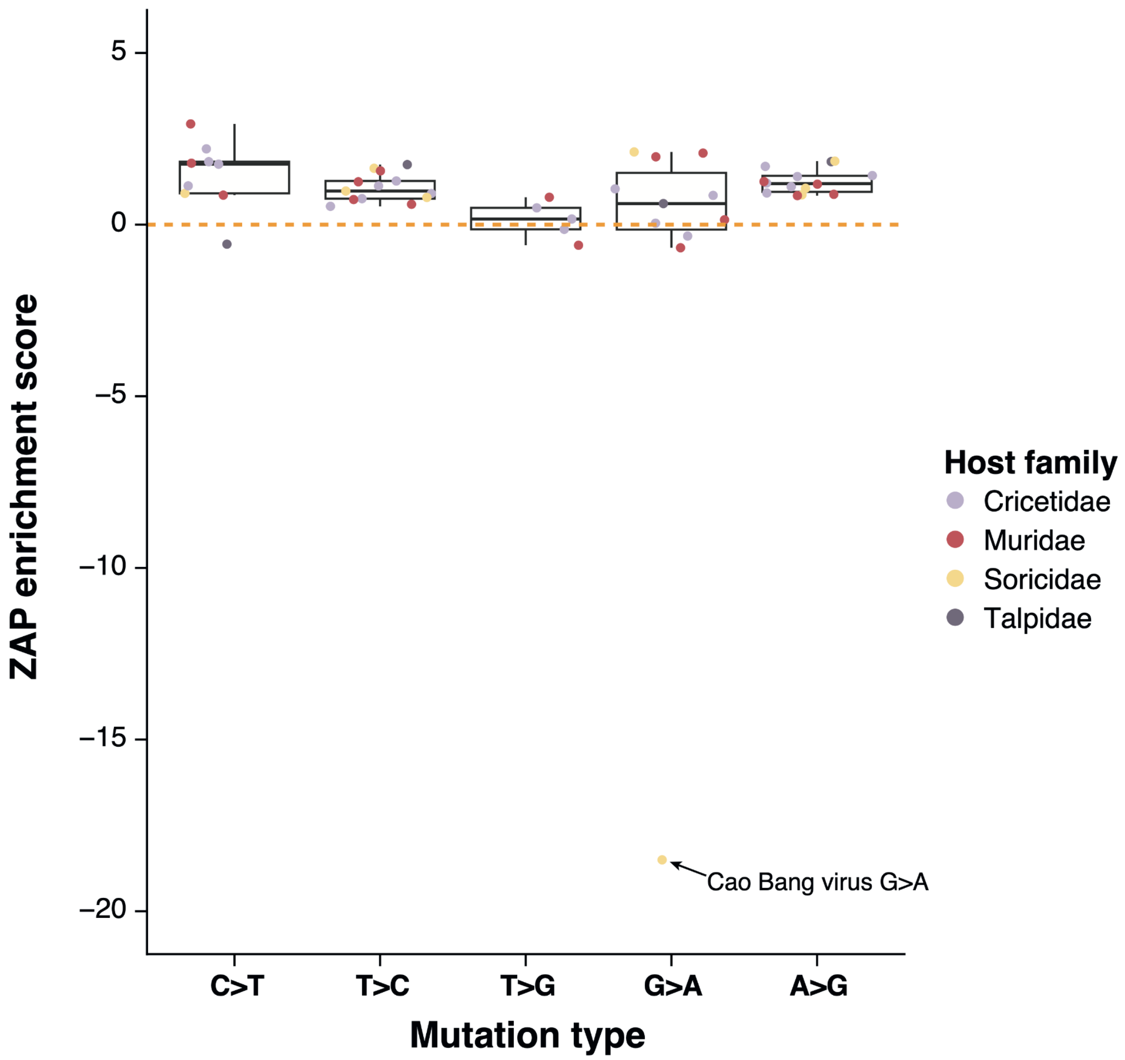


**Figure S7. Influence of host ZAP proteins on hantavirus mutagenesis including the Cao Bang virus G>T outlier.** We developed a ZAP enrichment score that calculates relative ZAP activity through the enrichment (for mutation types that remove ZAP contexts) or suppression (for mutation types that introduce ZAP contexts) of ZAP-associated contexts. This score was calculated for each mutation type in each virus that contains at least 160 mutations within the respective mutation type (see ***Methods***). The distributions of these scores are shown for each mutation type. T>A and A>T mutations are not included as they do not introduce or remove ZAP contexts. C>A, C>G, G>C, G>T and A>C are not shown as they contain a maximum of two viruses with at least 160 mutations. Cao Bang virus is an outlier in G>A mutations as it does not contain any G>T mutations in CG contexts despite 190 sampled G>A mutations.

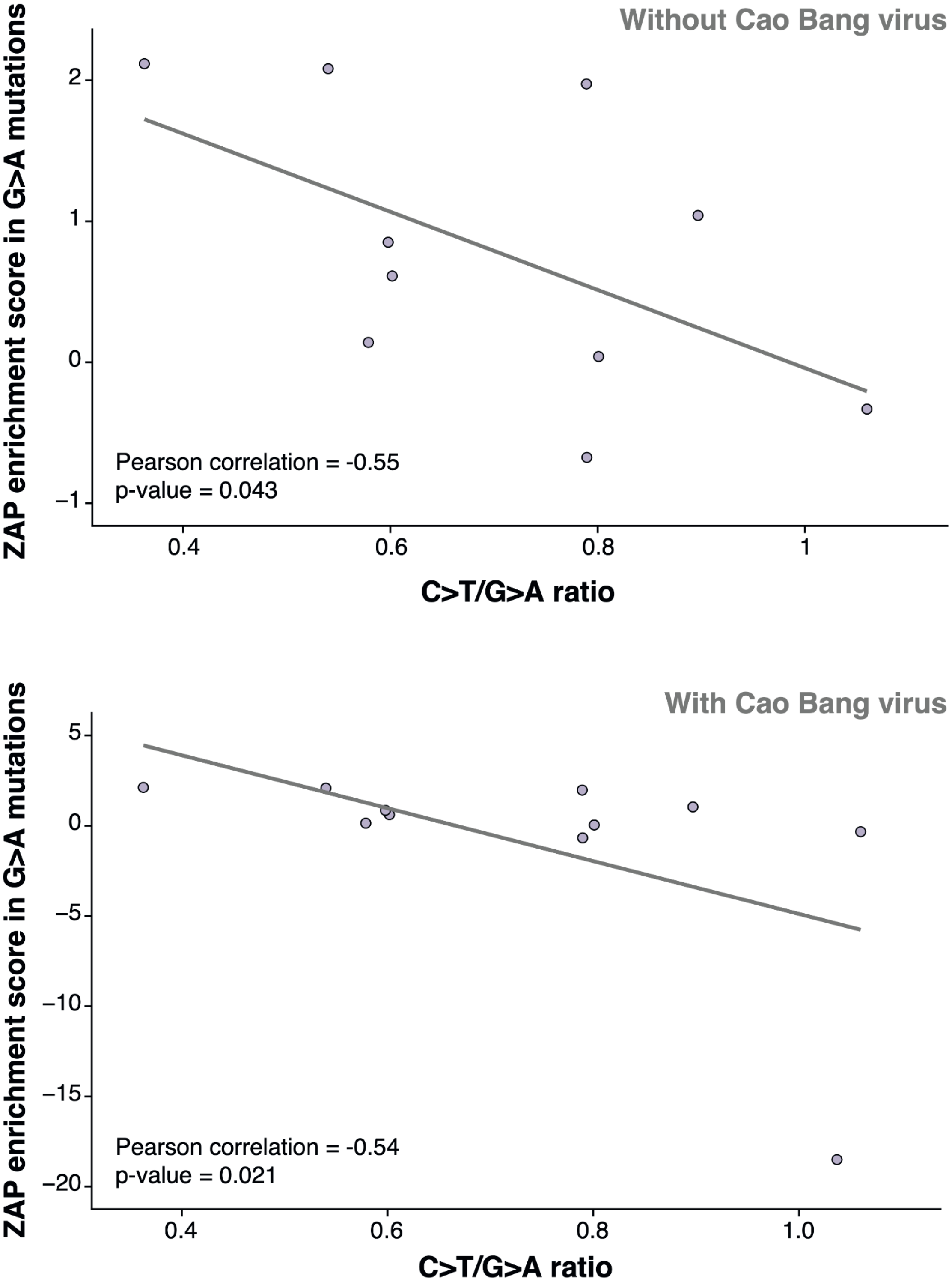


**Figure S8. Negative correlation between the ZAP enrichment in G>A mutations and the C>T/G>A ratio.** We calculated the ZAP enrichment score within G>A mutations for the 11 hantaviruses with at least 160 G>A mutations. This is plotted against the C>T/G>A symmetric mutation type ratio. All values are calculated within tree rescaled mutational spectra. The p-value on the correlation was calculated by comparing Pearson's r in the real data with that in 1000 randomisations of C>T/G>A ratios across viruses. Cao Bang virus is an outlier as it does not contain any G>A mutations in ZAP-associated contexts. We therefore carried out separate analyses excluding Cao Bang virus (top panel) and including Cao Bang virus (bottom panel).
